# Adapting antibacterial display to identify serum active macrocyclic peptide antibiotics

**DOI:** 10.1101/2023.07.28.550711

**Authors:** Justin R. Randall, Kyra E. Groover, Angela C. O’Donnell, Joseph M. Garza, T. Jeffrey Cole, Bryan W. Davies

**Affiliations:** Department of Molecular Biosciences, University of Texas at Austin, Austin, Texas 78712

## Abstract

The lack of available treatments for many antimicrobial resistant infections highlights the critical need for antibiotic discovery innovation. Peptides are an underappreciated antibiotic scaffold because they often suffer from proteolytic instability and toxicity towards human cells, making *in vivo* use challenging. To investigate sequence factors related to serum activity, we adapt an antibacterial display technology to screen a library of peptide macrocycles for antibacterial potential directly in human serum. We identify dozens of new macrocyclic peptide antibiotic sequences and find that serum activity within our library is influenced by peptide length, cationic charge, and the number of disulfide bonds present. Interestingly, an optimized version of our most active lead peptide permeates the outer membrane of gram-negative bacteria without strong inner membrane disruption and kills bacteria slowly while causing cell elongation. This contrasts with traditional cationic antimicrobial peptides, which kill rapidly via lysis of both bacterial membranes. Notably, this optimized variant is not toxic to mammalian cells and retains its function *in vivo*, suggesting therapeutic promise. Our results support the use of more physiologically relevant conditions when screening peptides for antimicrobial activity which retain *in vivo* functionality.

**Significance:** Traditional methods of natural antibiotic discovery are low throughput and cannot keep pace with the development of antimicrobial resistance. Synthetic peptide display technologies offer a high-throughput means of screening drug candidates, but rarely consider functionality beyond simple target binding and do not consider retention of function *in vivo*. Here, we adapt a function-based, antibacterial display technology to screen a large library of peptide macrocycles directly for bacterial growth inhibition in human serum. This screen identifies an optimized non-toxic macrocyclic peptide antibiotic retaining *in vivo* function, suggesting this advancement could increase clinical antibiotic discovery efficiency.

## Introduction

Antimicrobial resistant infections are now estimated to be a leading cause of death globally (1). While natural discovery of small molecule antibiotics continues to stagnate, a few novel macrocyclic peptide-based antibiotics like Teixobactin, Murepavadin, and Darobactin have recently been identified and show some clinical promise (2–4). These macrocyclic antimicrobial peptides (AMPs) can bind and inhibit essential cell envelope targets by adopting or mimicking β-sheet secondary structures (5–7). Other AMPs, like Colistin and many β-hairpin AMPs, are highly cationic and kill via disruption of the outer and inner bacterial cell membrane; however, they often also disrupt mammalian cell membranes causing toxicity (8–12). Though macrocyclic AMPs show some clinical promise, estimates suggest thousands if not millions of microbes need to be investigated to find one viable lead (13, 14). This highlights the need for innovation in the antibiotic discovery field (15).

Macrocyclic peptide drug discovery benefits from compatibility with both cell and cell-free display technologies, making them a promising group to explore synthetically (16). One *in vivo* function-based screening strategy, called surface localized antimicrobial display (SLAY), has been successfully used to identify macrocyclic peptides targeting the Gram-negative cell envelope (17). Most recently, this technique was used to identify a synthetic group of β-hairpin AMPs with enhanced bacterial specificity relative to natural members of the same class (18).

A major barrier to peptide antibiotics is their instability in biological fluids. To understand sequence features affecting this liability and to improve the efficiency with which macrocyclic peptide antibiotics with *in vivo* activity are discovered, we adapted SLAY to screen a peptide library based on natural β-hairpin AMP sequence features for inhibition of bacterial growth in human serum. This strategy led to the identification of sequence features associated with serum activity and a new lead peptide, SAP-26, which has unique killing activity, is non-toxic, and retains function *in vivo*.

## Results

### Previous AMPs identified using SLAY are serum inactive

Minimum inhibitory and bactericidal concentration assays (MICs and MBCs) are standard methods used to first examine antimicrobial potential. These assays are often performed against lab strains of bacteria grown in Mueller-Hinton (MH) media which is not representative of conditions found in the human body during infection. To determine how more *in vivo* relevant growth conditions impact SLAY identified AMP activity, we compared the MBC of multiple cationic AMPs against a laboratory strain of *E. coli* (W3110) and a clinically isolated strain (ATCC 25922) grown in both MH media and 100% human serum (HS). W3110 is a K12 derived strain and cannot grow in human serum and lacks O-antigen on its cell surface in contrast to 25922. MBCs were used instead of MICs because human serum was opaque, making bacterial growth difficult to observe by sight. We chose three cationic AMPs previously discovered using SLAY and four natural cationic AMPs, each with α-helical or β-hairpin secondary structure (18–24) (**Table S1**). All the SLAY identified AMPs showed no measurable activity against *E. coli* 25922 in serum while three of the four natural cationic AMPs retained some serum activity. Notably, the SLAY derived antimicrobials were identified in screens using a K12 laboratory strain of *E. coli* grown in Luria Broth (LB). Interestingly, both natural β-AMPs examined (Protegrin-1 and Tachyplesin-1), retained the most activity in human serum relative to MH, suggesting this is a promising class to examine under more *in vivo* relevant screening conditions.

### Design of a peptide library based on natural β-AMP sequence features

To identify synthetic AMPs with greater activity under biologically relevant conditions, we designed a 99,072-peptide library (BH) mimicking natural β-AMP attributes including residue frequency, length, charge, and potential number of disulfide bonds. To do this we used codon variation to represent the residue frequencies found in three regions of the β-AMP structure (tail, sheet, and loop) (**Fig. S1**). We potentiated multiple lengths via early stop codons and included the possibility of one to three cysteine pairs (**Fig. 1A, Fig. S1**). Inclusion of multiple arginine residues also allowed peptides in the library to range in charge from 4-12. We had twenty-four randomly selected peptides from this library (BHR) synthesized to determine their antimicrobial activity in both MH and HS (**Table S2**). Of the twenty-four peptides, 87.5% were active in MH media with a median MBC of 64 μg/ml and a range of 4 to 128 μg/ml. Only one of the twenty-four peptides (4%) retained any activity in human serum with an MBC of 128 μg/ml. This data confirmed the majority of the peptides in the library have antimicrobial activity, but lack strong activity in HS and are therefore ideal for testing our antimicrobial screening scheme in serum.

**Fig. 1:**
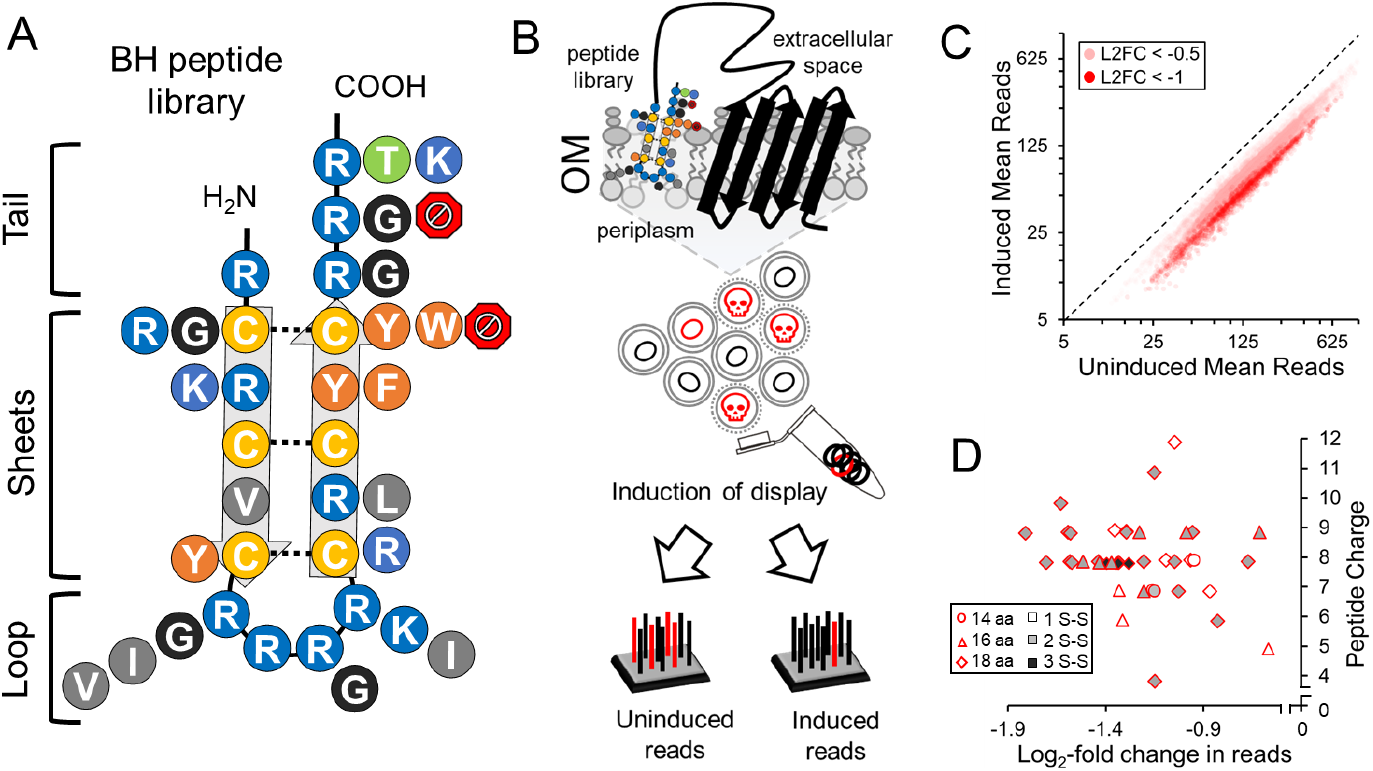
Synthetic discovery of serum active antimicrobial peptides. A) Diagram of the potential residues at each position of a peptide library based on natural β-AMP sequence frequencies. Potential disulfide bonds are indicated with a dotted line. Red octagons potentiate early stop codons. B) Diagram of how surface localized antimicrobial display functions. C) Uninduced versus induced mean reads for peptides in the library with a significant log_2_-fold change (L2FC) less than −0.5 or less than −1. D) Distribution of log_2_-fold change, charge, length and number of potential S-S bonds for select BHS peptides examined *in vitro*.

This library was cloned into a surface display plasmid and transformed into *E. coli* ATCC 25922 for testing.

### SLAY identifies serum active synthetic antimicrobial peptides

Surface localized antimicrobial display (SLAY) functions through the expression of a plasmid encoded fusion tethering a peptide to the outer membrane (**Fig. 1B**). During expression of the library, sequences with antimicrobial activity cause self-killing and therefore are reduced within the bacterial population. Differences between induced and uninduced plasmid copy number can be monitored and a log_2_-fold change generated by comparing induced and uninduced next-generation sequencing reads (**Fig. 1BC**). The resulting screen in *E. coli* 25922 identified 14,021 BH peptides with some potential for antimicrobial serum activity (log_2_-fold change ≤ −0.5, p-value ≤ 0.05). 1,598 of these peptides had strong antimicrobial potential (log_2_-fold change ≤ −1, p-value ≤ 0.05). We selected 41 peptides from this SLAY active group to examine *in vitro* (BHS) (**Fig. 1C, Table S2**). This group had a diverse range of charge (3.88-11.88), length (14-18 amino acids), and potential cysteine pairs (1-3) (**Fig. 1D**). All data relating to our SLAY screen can be found in supplemental data file 1.

The 41 selected SLAY active peptides were commercially synthesized and tested for their MBC in both MH and HS (BHS). The number of disulfide bonds present in each peptide was determined using high-resolution mass spectrometry paired with liquid chromatography (LC/MS) (**Table S3**). 97.6% of peptides killed E. coli 25922 in MH media while 34.1% remained active in human serum. This is over an eight-fold improvement in serum activity compared to the randomly selected peptides (BHR) (**Table S2**). BHS peptides with a length of 16 amino acids, 3 disulfide bonds, and charge of 6-8 were most likely to retain serum activity, suggesting these features improve potency (**Table 1**). No BHS peptides with a length less than 16 amino acids or charge greater than 9 retained any serum activity (**Table 1**).

**Table 1.**
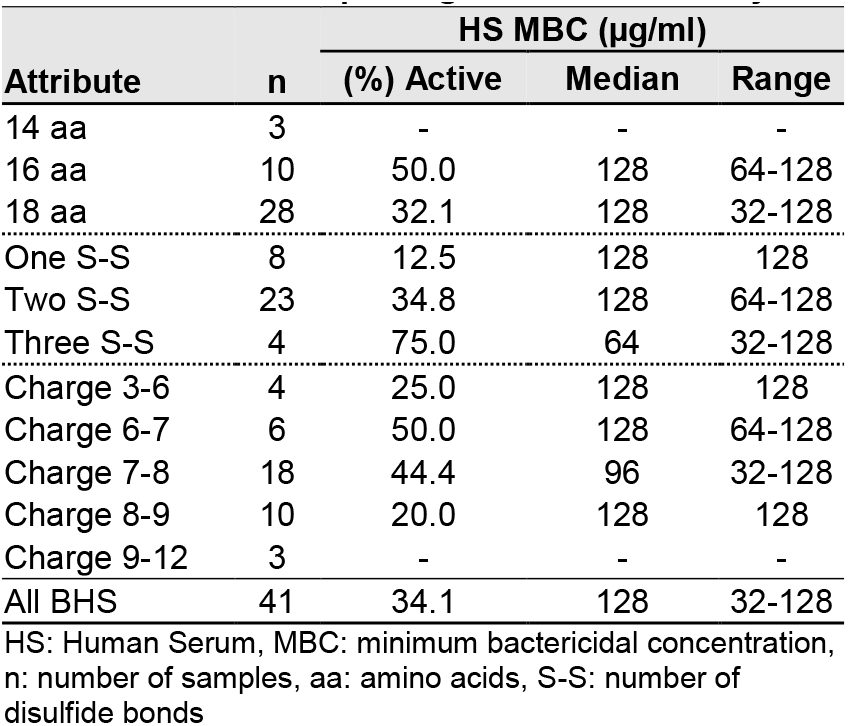
Attributes impacting BHS serum activity.

| Attribute | n | HS MBC ($\mu\text{g/ml}$) | | |
| --- | --- | --- | --- | --- |
|  |  | (%) Active | Median | Range |
| 14 aa | 3 | - | - | - |
| 16 aa | 10 | 50.0 | 128 | 64-128 |
| 18 aa | 28 | 32.1 | 128 | 32-128 |
| One S-S | 8 | 12.5 | 128 | 128 |
| Two S-S | 23 | 34.8 | 128 | 64-128 |
| Three S-S | 4 | 75.0 | 64 | 32-128 |
| Charge 3-6 | 4 | 25.0 | 128 | 128 |
| Charge 6-7 | 6 | 50.0 | 128 | 64-128 |
| Charge 7-8 | 18 | 44.4 | 96 | 32-128 |
| Charge 8-9 | 10 | 20.0 | 128 | 128 |
| Charge 9-12 | 3 | - | - | - |
| All BHS | 41 | 34.1 | 128 | 32-128 |

HS: Human Serum, MBC: minimum bactericidal concentration, n: number of samples, aa: amino acids, S-S: number of disulfide bonds

The most potent serum active peptide (BHS-18) was renamed SAP for <u>s</u>erum <u>a</u>ctive <u>p</u>eptide and further optimized to increase serum potency with the above data in mind (**Table S4**). Optimization included both canonical and non-canonical residue modifications, truncations, C-terminal amidation, N-terminal lipidation, and end to end cyclization. Truncation to 16 amino acids, C-terminal amidation, and use of 2,4-Diaminobutyric Acid (Dab) in the loop region all modestly improved serum potency while use of D-enantiomers, N-terminal lipidation, and end to end cyclization negatively impacted serum potency (**Table S4**). Ultimately, optimized variant SAP-26, with a four-fold improvement in serum potency over SAP, was selected for more detailed study.

### SAP-26 is an unstructured macrocyclic peptide

Many cationic AMPs demonstrate either β-hairpin or α-helical secondary structure upon interaction with bacterial membranes or membrane mimics like lipopolysaccharide (LPS) (25). To examine whether SAP-26 demonstrates a similar change in secondary structure we performed circular dichroism spectroscopy with and without the presence of 0.2 mg/ml LPS (**Fig. S2AB**). Surprisingly, SAP-26 showed an unstructured spectrum in buffer alone (minimum at ∼200 nm) and no significant shift in minimum upon addition of LPS. This was in contrast to known cationic AMPs Protegrin-1 (PG-1) and Cecropin P1 (CP1) whose minima shifted from unstructured to those consistent with β-sheet (∼218 nm) and α-helical (∼210 and 230 nm) structure respectively upon LPS addition (26) (**Fig. S2AB**). SAP-26 encodes six cystine residues. High resolution LC/MS of SAP-26 with and without a reducing agent, which reduces disulfide bonds, showed a shift of ∼6 Da in monoisotopic mass (M_mi_). This is consistent with the presence of three disulfide bonds (**Fig. S2CD**); however, we did observe a change in expected isotope distribution suggesting the SAP-26 molecular population contained a mixture of 1-3 disulfide bonds. Together, this data suggests that SAP-26 is macrocyclic but lacks strong secondary structure in solution with and without LPS present.

### SAP-26 activity is distinct from traditional cationic AMPs

Most cationic AMPs kill bacteria via disruption of the bacterial membrane so we compared SAP-26 activity to two known membrane lytic cationic AMPs (PG-1 and CP1) and two small molecule antibiotics (Kanamycin and Cefuroxime) which kill through ribosome and cell wall synthesis inhibition respectively (**Fig. 2A**). First, we performed two fluorescent membrane disruption assays using 1-N-phenylnaphthylamine (NPN) and propidium iodide (PI). Normally the outer membrane excludes hydrophobic molecules such as NPN, but when damaged, NPN can bind hydrophobic fatty acids within the membrane and fluoresce. PI fluoresces upon DNA binding but can only gain access to the cytoplasm if both the outer and inner membrane have been permeabilized. Antibiotics tested retained antimicrobial activity under the assay conditions used (**Fig. S3A**). Interestingly, SAP-26 demonstrated NPN fluorescence but not PI uptake after 30 minutes of treatment (**Fig. 2BC**). This was in contrast to PG-1 and CP1 which caused strong fluorescence of both NPN and PI. As expected Kanamycin, which can freely bypass cell membranes, showed no NPN fluorescence or PI uptake. SAP-26 began to show slight PI uptake at higher concentrations after two hours but this was still far less than was observed for PG-1 (**Fig. S3B**). This data supports SAP-26 quickly permeats the outer membrane without strong inner membrane disruption.

**Fig. 2:**
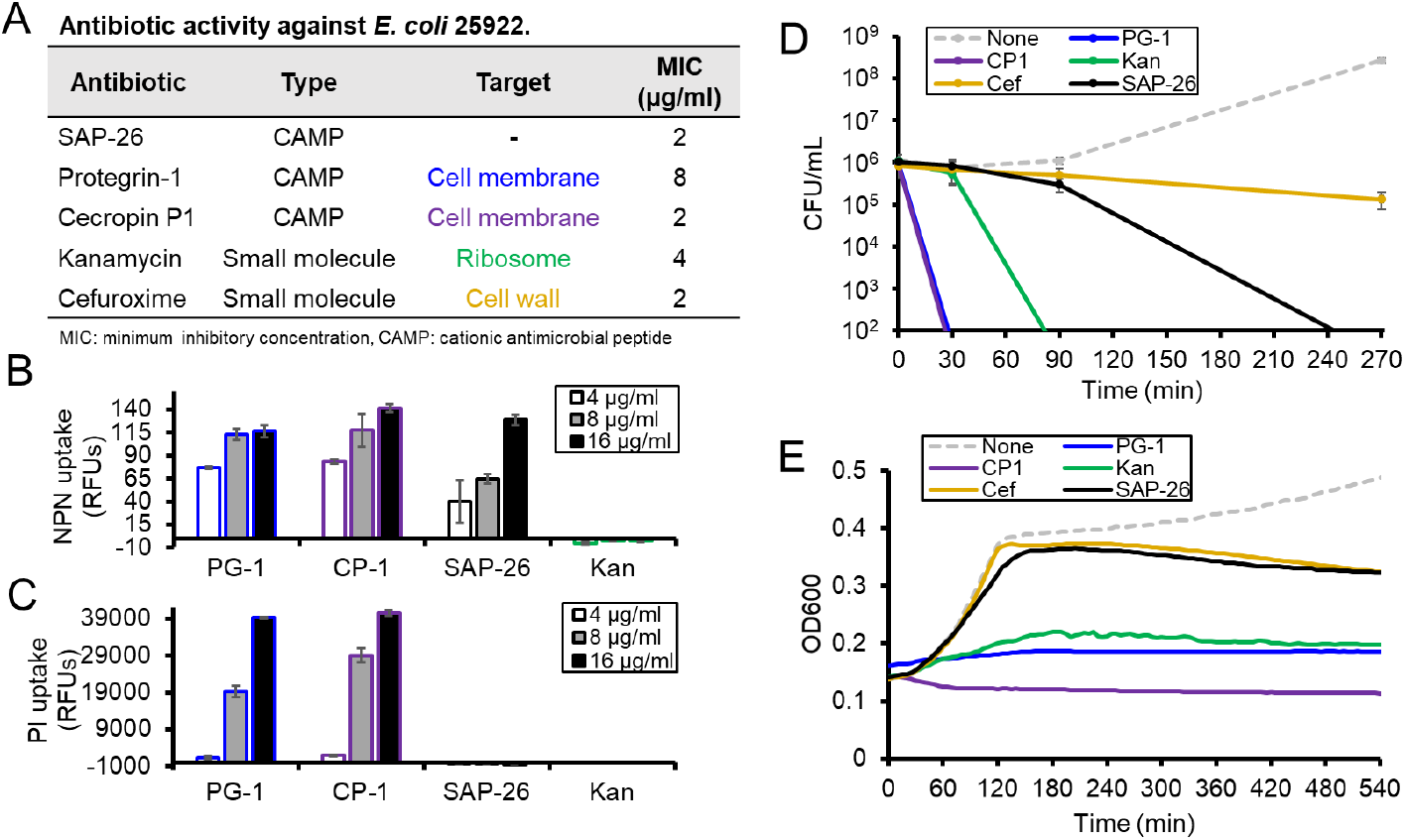
SAP-26 functions differently from traditional cationic AMPs. A) Table describing the structure and activity of various antibiotics. 1-N-phenylnaphthylamine (B) and propidium iodide (C) fluorescence of *E. coli* cells treated with various concentrations of antibiotic for 30 minutes. D) Kill curve of *E. coli* cells treated with antibiotic at 2x their listed MIC. E) Growth curve of *E. coli* cells treated with antibiotic at 8x their listed MIC. Listed MICs are the median of three replicates data points are an average of three replicates with error bars representing one standard deviation.

Membrane lytic cationic AMPs like PG-1 and CP1 kill cells rapidly, so we compared kill curves for *E. coli* 25922 treated with antibiotic at two-fold its MIC. SAP-26 did not kill cells rapidly like PG-1 and CP1, instead killing was delayed past that of Kanamycin but before Cefuroxime (**Fig. 2D**). A growth curve performed with each antibiotic added at eight-fold its MIC showed a similar result, with growth ultimately being inhibited later than CP1, PG-1, and Kanamycin but slightly before Cefuroxime (**Fig. 2E**). Together, this data further supports that SAP-26 is not killing through a rapid, inner membrane lytic mechanism like traditional cationic AMPs.

### SAP-26 is broad spectrum causes cell elongation and resistant mutants are difficult to isolate

To determine how SAP-26 kills bacterial cells, we first tested its spectrum of activity by determining its MIC against a panel of clinical and laboratory monoderm and diderm bacteria (**Table S5**). SAP-26 remained active against both groups and there was no clear evolutionary relationship between bacterial susceptibility. Interestingly a strain of *Corynebacterium striatum* was most susceptible while *Enterobacter cloacae* and *Klebsiella pneumoniae* strains examined were fully resistant. We also found SAP-26 activity was unaffected by the expression of mobile colistin resistance (mcr-1), suggesting it likely has a different mechanism of outer membrane disruption from the polymyxins (**Fig. S4A**).

Next, we attempted to identify SAP-26’s target through the isolation of resistant mutants via both plating and sub-inhibitory liquid passage in MH supplemented with SAP-26. To improve the likelihood of isolating a resistant mutant we used the *E. coli* Keio parent and a *mutS* deficient strain from the Keio collection with an increased mutation rate relative to wild type. Plating and serial passaging of both strains in the presence of SAP-26 resulted in no mutants with greater than two-fold resistance. In contrast, rifampicin resistant mutants were easily isolated via both methods (**Fig. S4B**). This data suggests that resistance to SAP-26 is not easily developed. To further evaluate whether SAP-26 may target an essential periplasmic protein, a pull-down was performed. Active N-terminal biotinylated SAP-26 was incubated with *E. coli* 25922 for two hours and then crosslinked using Lomant’s reagent. Cells were then lysed and SAP-26 was pulled-down with streptavidin beads. Biotin alone was used as a background binding control. Comparison of mass spectrometry between the two samples revealed only very low levels of protein binding specific to the SAP-26 sample and no obvious target. A full list of results are in supplementary data file 1.

Since attempted isolation of resistant mutants and pull-downs failed to identify a clear SAP-26 target, fluorescent microscopy with *E. coli* 25922 expressing cytoplasmic GFP was used to visualize possible changes in cell morphology. A small percentage (5.9%) of SAP-26 treated cells were severely elongated (cell length greater than two-fold the average untreated cell) (**Fig. 3A**, bottom panel). Even without this group of outliers we observed a modest but significant increase in mean cell length (p < 0.0001) (**Fig. 3A**). Interruptions in cell wall synthesis caused by beta-lactam antibiotics also cause cell elongation; however, knockout strains deficient in cell wall synthesis such as *dacA::kan, cpoB::kan*, and *mrcB::kan* showed no increased sensitivity to SAP-26, unlike beta-lactams, suggesting SAP-26 elongation is likely caused by a different process (27, 28) (**Fig. S4C**).

**Fig. 3:**
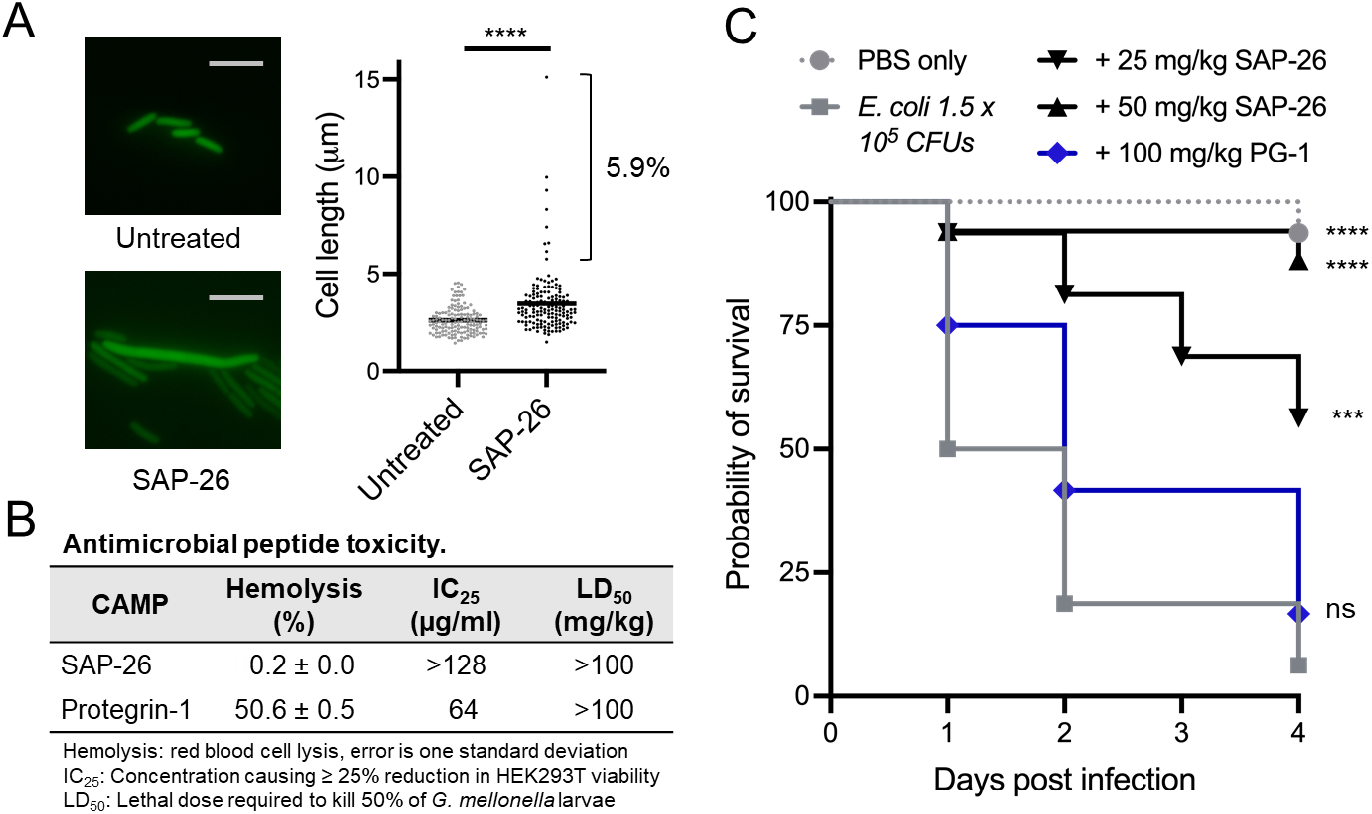
SAP-26 causes cell elongation, is non-toxic, and functions *in vivo*. A) Fluorescent microscopy images of *E. coli* 25922 cells expressing cytoplasmic GFP with and without treatment with SAP-26 (Scale = 5 μm). A scatter plot shows individual cell length and mean from both groups (n ≥ 148, **** p < 0.0001, Welch’s T-test). B) Table showing % hemolysis caused by 128 μg/ml AMP, IC_25_ for HEK293T tissue cultured cell, and LD_50_ for *G. mellonella* larvae. % Hemolysis error is one standard deviation of triplicate samples. C) Survival of *G. mellonella* larvae infected with *E. coli* 25922 +/-treatment with indicated concentrations of PG-1 or SAP-26. Significance was determined in relation to the untreated group (n ≥ 13, **** p < 0.0001, *** p = 0.0003, ns = not significant, log-rank test on Kaplan-Meier curves).

### SAP-26 is non-toxic, and remains active in vivo

Cationic AMPs commonly have mammalian cell toxicity making them difficult to develop as therapeutics. SAP and nearly all its derivatives, including SAP-26, show almost no hemolytic activity, in contrast to cationic AMPs like PG-1, a peptide included in the BH library design (**Fig. 3B, Table S4**). SAP-26 was also less toxic to embryonic kidney cells (HEK293T) than PG-1. Both AMPs LD_50_ against *G. mellonella* larvae was over 100 mg/kg. Peptide antibiotics also often lose functionality *in vivo* so we were curious if SAP-26 treatment could increase survival of *G. mellonella* larvae infected with 1.5 x 10^5^ *E. coli* 25922 cells (**Fig. 3C**). Treatment with a single dose of SAP-26 at either 25 or 50 mg/kg thirty minutes after infection significantly increased the probability of larvae survival relative to the untreated group (p = 0.0003 and p < 0.0001 respectively). Treatment with our highest dose of PG-1 (100 mg/kg) showed no significant increase in the probability of survival (p = 0.153). Together, this demonstrates that SAP-26 is both non-toxic and retains its antibacterial activity *in vivo*, in contrast to many cationic AMPs, suggesting it may be a promising lead for further mechanistic examination and clinical development.

## Discussion

Surface localized antimicrobial display is a promising synthetic method for peptide antibiotic discovery; however, it has thus far only identified membrane active peptides lacking activity in blood serum (**Table S1**) (17, 18). We adapted SLAY to function in conditions more representative of infection, identifying multiple macrocyclic antimicrobial peptides retaining activity in human serum (**Fig. 1**). Characteristics of many serum active peptides mimicked characteristics of potent natural β-AMPs including length (16-18), overall charge (6-8), and number of disulfide bonds (2-3) (**Table 1**). These same attributes are observed in Tachyplesin-1 and Protegrin-1 which were also shown to be serum active (Table S1). It appears that all AMPs examined lose activity in serum relative to broth; however, the extent of this loss in activity is variable and highly potent peptides are more likely to retain serum acticity. An increased number of disulfide bonds also correlated with increased serum activity which could be due to a reduction in proteolysis. This has been observed with other cationic AMPs (20). Reduced proteolysis could also explain the increased serum activity observed with SAP variants containing non-canonical residues and C-terminal amidation.

Our work strongly suggests that SAP-26 functions uniquely from traditional cationic AMPs like PG-1 and CP1, which kill via rapid outer and inner membrane disruption. We demonstrate SAP-26 permeates the outer membrane without strong inner membrane disruption and kills cells slowly (**Fig. 2)**. SAP-26 also causes cells to elongate, some drastically so (**Fig 3A**). These attributes are more similar to non-traditional cationic AMPs like Murepavadin, Thanatin, and JB-95 (29–31), which have been suggested to target essential cell envelope processes other than the inner membrane. Thanatin, which has been shown to permeate the outer membrane and target lipopolysaccharide transport, was included in the design of the beta-hairpin library design screened here. Additionally, SAP-26 retains activity against both monoderm and diderm bacteria (**Table S5**). Together, this data suggests that SAP-26 targets a still unresolved, broadly conserved, cell envelope process; however, a delayed lysis of the inner membrane cannot be completely ruled out as possible. Our inability to isolate SAP-26 resistant mutants or pull down a strong protein interactor implicates a substrate rather than an enzyme as the SAP-26 target. We hope to elucidate a more detailed mechanism of action in future studies.

Lastly, SAP-26 does not appear to have significant erythrocyte, kidney tissue, or *G. mellonella* larvae toxicity and retains its function *in vivo* against *G. mellonella* larvae infected with *E. coli* 25922 (**Fig. 3BC**). SAP-26 was especially active against a strain of *Corynebacterium striatum*, a growing nosocomial antimicrobial resistant pathogen (32). Unfortunately, the *C. striatum* strain examined here did not infect *G. mellonella* larvae so were unable to evaluate SAP-26 *in vivo* efficacy against this strain. Future therapeutic evaluation could be performed with clinically isolated antimicrobial resistant *C. striatum* strains in murine or other appropriate models of infection.

## Materials and Methods

### Growth inhibition and bactericidal assays

Stock peptides were diluted to 256 μg/ml in 350 μls of either MH or HS media and 100 μl was aliquoted in the top row of a polypropylene 96 well plate in triplicate. Peptides were then serial diluted two-fold down columns of the plate. Separately, each bacterial strain was grown overnight in either 5 ml of LB or Brain Heart Infusion (*Corynebacterium* and *Mycobacterium* strains only) media at 37 °C. Strains containing pDM1 and pDM1_mcr-1 were supplemented with 10 μg/ml tetracycline (33). Cells from overnight cultures were diluted to a concentration of ∼1 × 10^6^ cells/ml in either MH or HS and 50 μl were added to each well of the 96 well plate containing diluted peptide. For crude peptides only (BHR and BHS), acetic acid was added to 0.001% v/v and BSA to 0.02% w/v to improve solubility (34). Plates were incubated at 37 °C for 18-24 hours and examined by eye growth. For MBCs, five μl from each MIC plate was spotted on LB agar, dried, and incubated again overnight for observable growth. In cases where triplicate samples differed, the concentration supported by the median of the three replicates was reported.

### Peptide library cloning

Detailed methods for library creation have been previously reported (17, 35). Briefly, a library insert was generated by PCR using forward primer oJR557 with reverse primer oJR616 and 2x(NR)tether gBlock as the template (Table S6). The insert and the pMMBEH67_lpp_ompA vector were digested with KpnI and SalI and ligated overnight at 4°C using T4 ligase. The ligated library was cleaned and transformed into *E. coli* 25922 competent cells via electroporation for further analysis via SLAY.

### SLAY procedure

SLAY procedures have been detailed previously (17, 35). Briefly, *E. coli* 25922 frozen cells containing the BH plasmid library were diluted 1:1000 and recovered in 10 ml of HS supplemented with 75 μg/ml carbenicillin for 2 hours. The culture was then back diluted to OD 0.05 and three triplicate 5 ml cultures were set up in HS supplemented with 75 μg/ml carbenicillin. Triplicate reactions included: Uninduced (0 μM IPTG), and induced (100 μM IPTG). All triplicate cultures were then grown for 4 hours at 37°C. Plasmids from each triplicate culture were miniprepped and Illumina sequencing primers were used to produce an amplicon via PCR (Table S6). Amplicons were sent to Genewiz for next generation sequencing by Illumina MiSeq technology with 30% added Phi-X DNA.

### SLAY analysis

All possible codon combinations selected to generate the peptide sequences were encoded as nucleotide sequences for the reads to be mapped against. Flexbar v3.3.0 was used to trim the barcodes from the reads (36). The processed reads from flexbar were provided as input to Kallisto v0.46.1 to quantify the amount of reads that derived from each input codon sequence (37). The quantification file from Kallisto contained what would be akin to transcript level quantification where there are numerous potential codon combinations for a given peptide in the library. The R library tximport was used to take those values and transform them to peptide level read counts. The peptide level read counts were then provided as input to DESEQ2 to determine which peptides had significantly more read counts between the two libraries (38).

### Peptide Synthesis

All peptides used in this work were synthesized commercially by GenScript’s custom peptide synthesis service and analyzed by RP-HPLC and mass spectrometry to confirm molecular weight. For high purity peptides (> 90%) final concentration was determined using the molecular weight, A_205_ and A_205_ extinction coefficient. Final concentrations of these peptides were also adjusted for purity. A full list of all of the peptides used and their reported commercial purity can be found in supplemental data file 1.

### High-resolution LC/MS

The procedure used for LC/MS has been previously described (20). Briefly, peptides were diluted to 0.1 mM in PBS. Samples were separated by a C8 liquid chromatography column, and an Extracted Ion Chromatogram (EIC) was generated using an Agilent Technologies 6546 Accurate-Mass Q-TOF LC/MS instrument. Analysis was performed using Agilent MassHunter Qualitative software v10 and Agilent’s Isotope Distribution Calculator. For SAP-26 this was performed in triplicate with a representative spectrum shown. For crude peptides, this was performed once and the most abundant disulfide bond conformation was reported.

### Circular dichroism spectroscopy

Stock peptides were diluted in 10 mM potassium phosphate (pH 7.4) to 200 μg/ml in a volume of 200 μl. Samples were incubated 1-2 hours at room temperature and then analyzed using a Jasco-815 CD spectrometer with a 0.1 cm path-length quartz cuvette. The CD spectra were collected using far-UV spectra (190-250 nm) with background corrected for phosphate buffered saline alone. Ellipticity was then converted from mdeg to molar ellipticity. Reported spectra are an average of three separate spectra obtained from the same sample adjusted for molar concentration.

### NPN assays

A culture of E. coli 25922 was grown overnight in liquid media at 37°C. The following day, it was back diluted 1:25 in LB media and grown at 37°C until an OD600 of 0.5. The culture was then pelleted at 1000 g, washed once with PBS and resuspended to a final concentration of OD600 0.5. All experimental peptides were diluted in PBS to 128 μg/ml. Each sample was loaded in triplicate into a 96-well optical-bottom plate and serially diluted two-fold with final volumes of 50 μL in each well. 50 μL of the 40 μM NPN was then added to each well, followed by 100 μL of the bacterial cell suspension. After 30 minutes the plate was transferred to a BioTek Synergy LX multi-mode reader for fluorescent measurements. Triplicate samples were normalized to non-treated wells and presented as a mean ± one standard deviation (n=3)

### Propidium iodide assays

Propidium iodide uptake was measured for *E. coli* 25922 as previously described (20). Briefly, single colonies from overnight growth on LB were inoculated into Mueller Hinton broth and grown to mid-log phase. Cells were then washed twice with PBS + 50 mM glucose and resuspended to an OD = 0.1 in PBS + 50 mM glucose. Propidium iodide was added at a concentration of 10 μg/ml and 50 μL of the propidium iodide cell mixture was quickly added to a pre-prepared 96-well plate (NUNC black-walled clear bottom plate) that contained 50 μl of various peptide concentrations. The plate was then allowed to incubate for 30 minutes in the dark at 37°C and was then read using the Biotek Synergy LX plate reader with the fluorescence red filter cube. Triplicate samples were normalized to non-treated wells and presented as a mean ± one standard deviation (n = 3).

### Bacterial kill curves

*E. coli* 25922 cells were back diluted from an overnight LB into MH to an OD of 0.001. Peptides stock solutions were diluted in MH to four times their reported MH MIC. 100 μl of diluted cells were combined with 100 μl of diluted peptide stock in triplicate. 10 μl of each sample was then serially diluted ten-fold at the indicated time points and plated on LB agar. CFUs were counted after overnight growth at 30°C. Error bars represent one standard deviation of triplicate samples. No peptide addition was used as a negative control.

### Bacterial growth curves

*E. coli* 25922 cells were back diluted 50-fold from an overnight LB culture in MH and grown to an OD of 0.2. Peptides stock solutions were diluted in MH to 16 times their reported MH MIC. 100 μl of diluted cells were combined with 100 μl of diluted peptide stock into a clear flat bottom 96 well plate in triplicate. Cell were grown at 37°C shaking for 9 hours in a BioTek Logphase600 microbiology plate reader with the OD taken every 5 minutes. Growth curves are reported as the mean representing triplicate samples.

### Microscopy

*E. coli* 25922 cells were freshly transformed with the pUltraGFP plasmid (39) and grown in MH media to logarithmic phase growth (OD = 0.3-0.5). Cells were then concentrated four-fold via centrifugation and 5 μl spotted on a microscope slide with a MH pad containing 1.1% agarose with or without 16 μg/ml SAP-26 and covered. Slides were incubated at room temperature for 30-60 minutes and then imaged with a Nikon Eclipse TE2000-U microscope with 100x objective and GFP filter. Cell length was calculated for individual cells using NIS-Element AR software and the mean was reported. Significance was determined by a Welch T-test.

### Generation of phylogenetic tree

Full genomes were downloaded from the NCBI database (www.ncbi.nlm.nih.gov). Phylogenetic reconstruction was generated using PhyloPhlAn version 3.0.67 (40). Within the pipeline, Diamond was used for genome mapping, MAFFT was used to generate multi-sequence alignment, and IQ-Tree was specified for phylogeny building (41, 42). Output from PhyloPhlAn was visualized as a cladogram using the ggtree (v 3.6.2) package in R (v 4.2.2) (43).

*SAP-26 Pull-Down. E. coli* 25922 cells were grown overnight in 20 ml of MH media, pelleted, washed, and concentrated to an OD = 10 in 1.8 ml of MH. Cell were split into two 900 ul samples and either SAP-26 or Biotin was added to cells at 542 μM. Cells were incubated for two hours at 37 °C crosslinked using Lomant’s reagent and 1x protease inhibitors and 0.1% triton-X100 was added. Cell were lysed via sonication, debris was pelleted, and the supernatant containing was removed and treated with nuclease at 37 °C for one hour. 20 μl of washed Strep-Tactin beads were then added to each sample and incubated for 1 hour at 4 °C. Each sample was washed 2x and then eluted in 50 ul of PBS by boiling. Samples were then submitted to the UT Proteomics core for protein identification analysis. Results show total spectral counts for proteins found specifically in the SAP-26 containing sample only (with exception of streptavidin).

### Hemolysis assays

Single donor human red blood cells were washed in PBS and adjusted to a concentration of 1 x 10^9^ cells/ml. Each peptide was added in triplicate to 200 μl of cells at a concentration of 128 μg/ml in a 96 well polypropylene plate. PBS alone and 1% triton-X100 were used for background normalization and 100% hemolysis respectively. Plates were incubated for 3 hours at 37 ºC. Following incubation samples were centrifuged at 800 g for 20 min and 100 μl of supernatant was transferred to a flat bottom 96 well plate. Percent hemolysis for each sample was determined by normalizing the absorbance at 540 nm for each sample to the average background and dividing by the average absorbance for 1% Triton X-100 (100% hemolysis). Error bars represent one standard deviation of triplicate samples.

### Cytotoxicity

Adherent HEK293T cells were grown in DMEM with the addition of 10% FBS with penicillin and streptomycin at 37°C with 5% CO_2_. For the assay 5,000 cells were seeded per well in culture medium and allowed to grow for 24 hours. After 24 hours the media was replaced with fresh media. Peptides were prepared on a separate plate at 10X the concentration and serially diluted two-fold starting with 1,280 μg/ml. Cells and peptides were incubated for 48 hours at 37°C with 5% CO_2_. After the media was removed and serum free DMEM with 0.5mg/mL MTT was added to the cells. The plate was incubated at 37°C for four hours. The media was removed and the MTT crystals were dissolved with MTT solvent (isopropanol, 0.1% igepal, and 4mM HCl). The plate was shaken for 15 minutes and then read at 570 nm. Percent viability was determined by dividing absorbance by the average of the untreated wells which were considered 100% viable. IC_25_ was reported as the concentration of peptide necessary to cause at least 25% reduction in viability.

### Galleria mellonella infection model

Live *Galleria mellonella* larvae were purchased from DBD Pet. Larvae appearing fully healthy (no apparent melanization) and weighing between 200-300 mg were used for survival assays. Larvae were first injected into their second left proleg with 10 μl of PBS or PBS containing 1 x 10^6^ *E. coli* 25922 cells grown to an OD of 0.5 and washed in PBS. Injections were performed using a Hamilton 250 μl pipet equipped with an auto-dispenser and 30-gauge BD needle. Larvae then received a second 10 μl injection into their second right proleg containing PBS or PBS containing cationic AMP (SAP-26, PG-1, or CP1) diluted to 100 mg/kg. Larvae groups were then incubated at 37 °C for 72 hours and observed for death. Larvae were determined to be dead if they were unable to right themselves after being placed on their back. Significance was determined by comparing Kaplan-Meier survival curves to untreated using a log-rank test (n ≥ 13 per sample). For LD_50_ a single injection containing a 100 mg/kg dose of AMP was performed and worms monitored for death after four days (n = 10).

## Acknowledgements

We would like to thank the Targeted Therapeutic Drug Discovery and Development Program at the University of Texas for access to circular dichroism training and equipment, the Mass Spectrometry Facility at The University of Texas at Austin for help with high-resolution LC/MS, and the UT Austin Center for Biomedical Research Support Biological Mass Spectrometry Facility (RRID:SCR_021728) for pull-down protein identification. Also, thanks to Nancy Moran’s lab for the use of their microscope and Despoina Mavridou’s lab for sharing mcr-1 and GFP expressing plasmids.

## Author Contributions

Conceptualization: JR, BD

Methodology: JR, BD

Investigation: JR, KG, AO, JG, JC

Visualization: JR, AO

Funding acquisition: BD

Project administration: JR, BD

Supervision: JR, BD

Writing – original draft: JR

Writing – review & editing: JR, BD

## Data Sharing Plan

Raw sequencing data from the SLAY experiment will be uploaded to the SRA database. All other data are available in the main text or the supplementary materials.

