## Supplemental Figures and Tables for "Adapting antibacterial display to identify serum active macrocyclic peptide antibiotics"

**This PDF includes:**

Supplemental Figures S1-4
Supplemental Tables S1-S6

A

| KNOWN B-L-B AMPS | | #AA | Charged AAs |
|---|---|---|---|
| POLYPHEMUSIN-1 | RRWCFRVCYRGFCYRKCR | 18 | 7 |
| TIGERININ-1 | FCTMIPIPRCY | 11 | 1 |
| BECTENECIN | RLCRIVVIRVCR | 12 | 4 |
| THANATIN | GSKKPVPIIYCNRRTGKCQRM | 21 | 6 |
| ARENICIN-1 | RWCVYAYVRIRGVLVRYRRCW | 21 | 6 |
| TACHYPLESIN-1 | KWCFRVCYRGICYRR | 17 | 6 |
| GOMESIN | CRRLCYKQRCVTYCRGR | 18 | 6 |
| ANDROCTONIN | RSVCRQIKICRRRGGCYYKCTNRPY | 25 | 8 |
| PROTEGRIN-1 | RGGRLCYCRRRFCVCVGR | 18 | 6 |
| DEFENSIN-1 | GFCRCLCRRGVCRCICTR | 17 | 5 |
| | Averages | 17.9 | 5.6 |
| | Library | 14,16,18 | 2-13 |

B

| AA | Frequency | | AA | Frequency | | AA | Frequency | |
|---|---|---|---|---|---|---|---|---|
| Tails | Natural | Library | Sheets | Natural | Library | Loop | Natural | Library |
| R | 0.31 | 0.40 | C | 0.36 | 0.30 | R | 0.32 | 0.52 |
| G | 0.10 | 0.17 | R | 0.20 | 0.16 | G | 0.15 | 0.19 |
| K | 0.07 | 0.10 | Y | 0.09 | 0.10 | I | 0.11 | 0.15 |
| V | 0.07 | - | V | 0.09 | 0.10 | V | 0.09 | 0.06 |
| F | 0.07 | 0.06 | K | 0.05 | 0.07 | K | 0.06 | 0.08 |
| Y | 0.07 | 0.04 | I | 0.04 | 0.05 | Y | 0.06 | - |
| W | 0.07 | 0.04 | L | 0.03 | 0.04 | F | 0.04 | - |
| P | 0.05 | - | T | 0.03 | 0.03 | L | 0.04 | - |
| T | 0.03 | 0.04 | F | 0.02 | 0.04 | M | 0.04 | - |
| I | 0.03 | - | W | 0.02 | 0.02 | P | 0.04 | - |
| S | 0.03 | - | A | 0.02 | - | Q | 0.02 | - |
| L | 0.03 | - | Q | 0.01 | - | T | 0.02 | - |
| N | 0.02 | - | P | 0.01 | - | | | |
| A | 0.02 | - | S | 0.01 | - | | | |
| M | 0.02 | - | G | 0.01 | - | | | |

C

| Position | 0 | 1 | 2 | 3 | 4 | 5 | 6 | 7 | 8 | 9 | 10 | 11 | 12 | 13 | 14 | 15 | 16 | 17 |
|---|---|---|---|---|---|---|---|---|---|---|---|---|---|---|---|---|---|---|
| AA(s) | R | CGR | RK | C | V | CY | RIGV | R | GR | RKI | CR | RL | C | YF | CYW[S] | RG | RG[S] | RTK |
| # Possible | 1 | 3 | 2 | 1 | 1 | 2 | 4 | 1 | 2 | 3 | 2 | 2 | 1 | 2 | 3 | 2 | 2 | 3 |
| Charge | 1 | 0.33 | 1 | | | | 0.25 | 1 | 0.5 | 0.66 | 0.5 | 0.5 | | | | 0.5 | 0.5 | 0.66 |
| Codon | **AGG** | **BGC**<br>CGC<br>TGC<br>GGC | **ARA**<br>AAA<br>AGA | **TGC** | **GTT** | **TRT**<br>TGT<br>TAT | **RKA**<br>AGA<br>ATA<br>GGA<br>GTA | **AGG** | **RGA**<br>AGA<br>GGA | **ADA**<br>AGA<br>AAA<br>ATA | **YGC**<br>TGC<br>CGC | **CKA**<br>CGA<br>CTA | **TGC**<br>TGC | **TWC**<br>TAC<br>TTC | **TRK**<br>TAG<br>TAT<br>TGT<br>TGG | **RGA**<br>GGA<br>AGA | **BGA**<br>CGA<br>TGA<br>GGA | **AVA**<br>AGA<br>AAA<br>ACA |

| | 14aa | 16aa | 18aa | | |
|---|---|---|---|---|---|
| Length | 14aa | 16aa | 18aa | | |
| Average charge | 5.74 | 6.24 | 7.4 | | |
| Unique peptides | 2304 | 13824 | 82944 | Total | 99,072 |

Reverse primer TBT TCV TCY MYA GWA GCA TMG GCR THT TCY CCT TMY AYA AAC GCA TYT GCV CCT

**Figure S1: Design of a peptide library based on natural beta-AMP residue frequency.** A) Amino acid sequences for ten natural beta-AMP sequences. B) individual residue frequencies found in the tail, sheet, and loop regions of the ten-natural beta-AMPs and the BH peptide library. C) Codons used to cod for amino acids at each position of the BH peptide library.

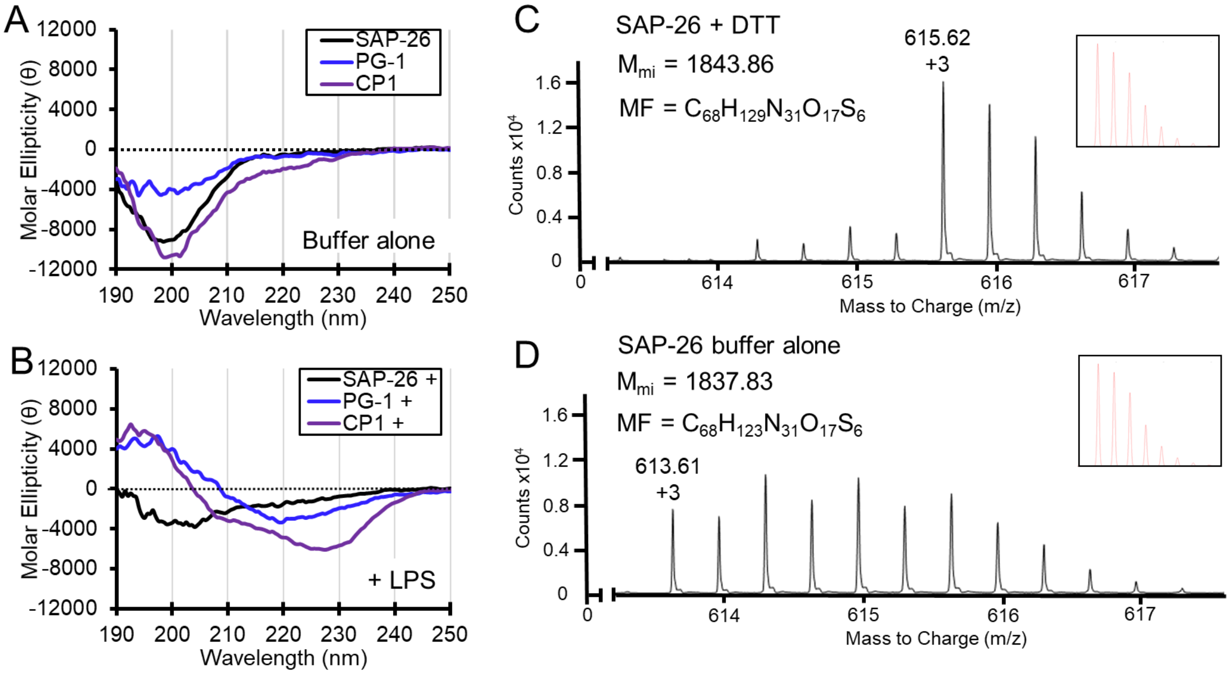


**Figure S2: SAP-26 is an unstructured peptide macrocycle.** Circular dichroism spectra of SAP-26, Protegrin-1 (PG-1), and Cecropin P1 (CP1) in buffer (A) with 0.2 mg/ml LPS (B). Data represents the mean of three technical replicates. High-resolution mass spectrometry of SAP-26 showing individual isotopes with 10 mM DTT (C) or in buffer alone (D). Mass to charge and charge state are highlighted for the peak corresponding to the monoisotopic mass ($M_{mi}$). The calculated $M_{mi}$ and molecular formula (MF) are shown and expected isotope distribution is inset.

A

***E. coli* 25922 antibiotic killing in PBS glucose**

| Antibiotic | MBC (µg/ml) |
|---|---|
| SAP-26 | 1 |
| Protegrin-1 | 2 |
| Cecropin P1 | 2 |
| Kanamycin | 1 |

MBC: minimum bactericidal concentration,

B

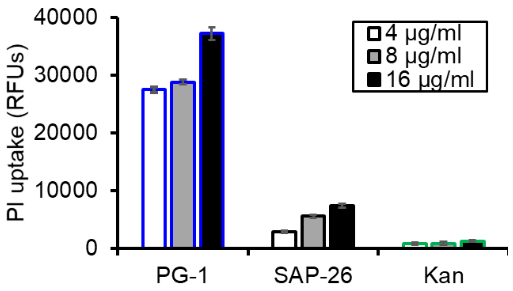


**Figure S3: Comparing SAP-26 activity to other antibiotics.** A) Table showing the MBC of antibiotics used in Fig. 3 in PBS supplemented with 50 mM glucose. B) PI fluorescence at various concentrations for cells treated with Protegrin-1 (PG-1), SAP-26, and Kanamycin (Kan) after two hours.

A

**Effect of colistin resistance on SAP-26**

| Strain | SAP-26 MIC (µg/ml) | Colistin MIC (µg/ml) |
|---|---|---|
| E. coli W3110 | 1 | 0.5 |
| + pDM1_empty | 1 | 0.5 |
| + pDM1_mcr-1 | 1 | 8 |

MH: Mueller-Hinton, MIC: minimum bactericidal concentration

B

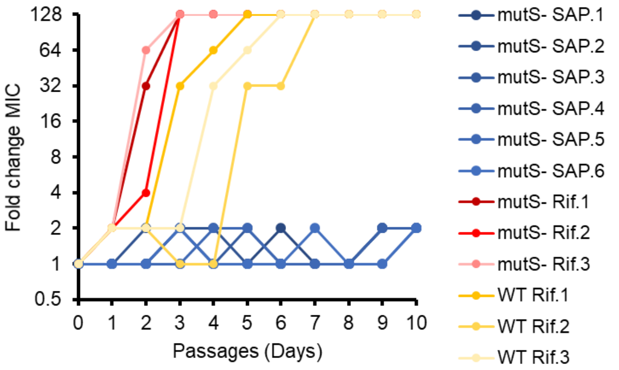


C

**Beta-lactam sensitive knockout MICs.**

| Antibiotic | Keio WT MIC (ug/ml) | *dacA::kan* MIC (ug/ml) | *cpoB::kan* MIC (ug/ml) | *mrcB::kan* MIC (ug/ml) |
|---|---|---|---|---|
| SAP-26 | 1 | 1 | 2 | 1 |
| Protegrin-1 | 8 | 8 | 8 | 8 |
| Carb | 8 | 1 | ≤ 1 | 2 |
| Cefuroxime | 8 | 2 | 1 | 1 |

MIC: minimum inhibitory concentration, Carb: carbenicillin

**Figure S4: SAP-26 resistance and cell wall deficient insensitivity.** A) Table showing SAP-26 activity against E. coli W3110 containing an IPTG inducible empty vector or mcr-1 expressing plasmids. IPTG was added to 0.5 mM. B) Graph showing the Fold-change in MIC for independent isolates of *E. coli* K12 (WT) and *mutS* deficient (mutS-) strains during ten days of serial passage in sub inhibitory concentrations of SAP-26 (SAP) or Rifampin (Rif). C) Table showing the MIC of antibiotics against strains deficient in cell wall synthesis.

**Table S1. Effect of bacterial strain and media conditions on CAMP activity.**

| | | | *E. coli* W3110 MBC (µg/ml) | *E. coli* 25922 MBC (µg/ml) | |
|---|---|---|---|---|---|
| **CAMP** | **Source** | **Structure** | **Mueller-Hinton** | **Mueller-Hinton** | **Human Serum** |
| PepC | SLAY | α-helical | 32 | 64 | >256 |
| Symbah-1 | SLAY | β-hairpin | 16 | 32 | >256 |
| SySA-5 | SLAY | β-hairpin | 16 | 64 | >256 |
| Melittin | Nature | α-helical | 16 | 8 | >256 |
| Cecropin P1 | Nature | α-helical | 2 | 2 | 32 |
| Protegrin-1 | Nature | β-hairpin | 4 | 8 | 32 |
| Tachyplesin-1 | Nature | β-hairpin | 4 | 8 | 16 |

CAMP: cationic antimicrobial peptide, MBC: minimum bactericidal concentration

**Table S2. Characteristics of randomly selected BH library peptides.**

| Name | Sequence | MBC MH | MBC HS |
|---|---|---|---|
| BHR-1 | RCKCVYRRRIRRCFCRRR | 8 | >128 |
| BHR-2 | RGKCVYGRRRRRCYWRGR | 64 | >128 |
| BHR-3 | RRKCVCGRRICRCYCG | >128 | >128 |
| BHR-4 | RRKCVYIRRIRRCYWRGT | 8 | 128 |
| BHR-5 | RRRCVYRRRKCRCYWGGT | 64 | >128 |
| BHR-6 | RGKCVCIRGRRLCYYR | 32 | >128 |
| BHR-7 | RRRCVYIRGICRCFYGGR | 32 | >128 |
| BHR-8 | RRKCVCGRRKRLCYYRRK | 64 | >128 |
| BHR-9 | RGRCVCRRGICLCYWRRK | 4 | >128 |
| BHR-10 | RCRCVCVRGKRRCFWGGK | 128 | >128 |
| BHR-11 | RGRCVYRRGICLCFYGRR | 32 | >128 |
| BHR-12 | RGKCVYIRGICLCYYGGT | >128 | >128 |
| BHR-13 | RRKCVYRRRKCLCYWR | 64 | >128 |
| BHR-14 | RCRCVCGRGRRRCYCRRK | 64 | >128 |
| BHR-15 | RCKCVYVRGKCRCFCGGR | 64 | >128 |
| BHR-16 | RRKCVCIRRICLCYCGGR | 128 | >128 |
| BHR-17 | RCKCVYVRRRCRCYYG | 64 | >128 |
| BHR-18 | RRRCVCVRRRCLCFYRRR | 128 | >128 |
| BHR-19 | RGKCVCVRRIRLCFCG | 4 | >128 |
| BHR-20 | RGKCVCRRGKRRCYYGRR | 128 | >128 |
| BHR-21 | RGRCVCIRGRRLCYWGGK | >128 | >128 |
| BHR-22 | RRKCVYIRGKRRCYWGRK | 128 | >128 |
| BHR-23 | RCKCVCIRGRCRCFWRGK | 32 | >128 |
| BHR-24 | RGKCVCIRRKRRCYCRRK | 128 | >128 |
| **% Active** | – | **87.5** | **4.0** |
| **Median MBC** | – | **64** | **128** |

MBC: minimum bactericidal concentration, MH: Mueller-Hinton;
HS: Human Serum

### Table S3: SLAY active BH peptide biochemical characteristics.

| Name | Sequence | MH MBC | HS MBC | L2FC | Length | #S-S | Charge |
|---|---|---|---|---|---|---|---|
| BHS-1 | RRRCVYRRRRRRCFCRRR | 64 | >128 | -1.05 | 18 | 1 | 11.88 |
| BHS-2 | RCKCVYGRGICLCFCGGR | 128 | >128 | -1.15 | 18 | nd | 3.82 |
| BHS-3 | RRRCVCVRRRRRCFCRRR | 64 | >128 | -1.15 | 18 | 2 | 10.85 |
| BHS-4 | RGRCVYVRRIRLCYYG | 16 | >128 | -0.57 | 16 | 1 | 4.91 |
| BHS-5 | RRRCVYVRGICLCYCRRR | 16 | 128 | -0.87 | 18 | nd | 6.85 |
| BHS-6 | RGRCVYVRGKRLCYWRRR | 8 | >128 | -0.97 | 18 | 1 | 7.91 |
| BHS-7 | RGKCVCIRRICRCYWGRT | 2 | >128 | -0.83 | 18 | 2 | 5.85 |
| BHS-8 | RGKCVCRRGRRLCFCGGK | >128 | >128 | -1.03 | 18 | 2 | 6.85 |
| BHS-9 | RGRCVCVRGKCRCYWRRK | 8 | 128 | -1.05 | 18 | 2 | 7.85 |
| BHS-10 | RGRCVCRRRRCRCY | 128 | >128 | -1.15 | 14 | 2 | 6.85 |
| BHS-11 | RCRCVYGRGRRLCFWR | 4 | 128 | -1.31 | 16 | 1 | 5.88 |
| BHS-12 | RGKCVYRRGKCRCFWR | 32 | 128 | -1.33 | 16 | nd | 6.88 |
| BHS-13 | RCRCVCRRRKCRCFWR | 8 | >128 | -1.37 | 16 | 2 | 7.82 |
| BHS-14 | RRRCVCIRRKRLCFCG | 2 | 64 | -1.21 | 16 | 2 | 6.85 |
| BHS-15 | RCRCVYVRRKRRCYYRGR | 64 | >128 | -1.29 | 18 | 1 | 8.88 |
| BHS-16 | RCKCVYIRRRRLCYCRGR | 2 | 64 | -1.44 | 18 | 2 | 7.85 |
| BHS-17 | RCRCVCIRRRRRCFCGGK | 32 | >128 | -1.57 | 18 | 2 | 7.82 |
| BHS-18 | RCRCVCRRRKCRCYCGRT | 1 | 32 | -1.28 | 18 | 3 | 7.79 |
| BHS-19 | RCRCVYIRRRRRCF | 64 | >128 | -1.16 | 14 | 1 | 6.88 |
| BHS-20 | RGKCVCRRRRCRCYWR | 4 | 128 | -1.52 | 16 | 2 | 7.85 |
| BHS-21 | RGRCVCIRRRCRCYYRGR | 8 | 128 | -1.58 | 18 | 2 | 7.85 |
| BHS-22 | RGRCVCRRRKCRCYCGGR | 64 | >128 | -1.7 | 18 | nd | 7.82 |
| BHS-23 | RCKCVCRRRRCRCYWRGR | 16 | >128 | -1.58 | 18 | nd | 8.82 |
| BHS-24 | RCRCVCRRGKRRCFCRRR | 128 | >128 | -1.63 | 18 | 2 | 9.82 |
| BHS-25 | RCKCVCIRRKCRCFCRGK | 8 | >128 | -1.34 | 18 | 3 | 7.79 |
| BHS-26 | RCKCVCIRRKCRCYCRGK | 8 | 128 | -1.4 | 18 | 3 | 7.79 |
| BHS-27 | RGRCVCRRRRRRCF | 128 | >128 | -0.95 | 14 | nd | 7.88 |
| BHS-28 | RCRCVYRRRRRRCFCR | 32 | >128 | -0.99 | 16 | 2 | 8.85 |
| BHS-29 | RCRCVYRRRRRRCYCR | 16 | 128 | -0.61 | 16 | 2 | 8.85 |
| BHS-30 | RGRCVCGRRRCLCYYRRR | 16 | >128 | -0.67 | 18 | 2 | 7.85 |
| BHS-31 | RCRCVCRRRRCRCFWR | 16 | >128 | -1.43 | 16 | 2 | 7.82 |
| BHS-32 | RCKCVYRRRRRRCFCR | 64 | >128 | -1.23 | 16 | 2 | 8.85 |
| BHS-33 | RRKCVYIRRRRRCYWGGR | 32 | >128 | -1.35 | 18 | 1 | 8.91 |
| BHS-34 | RRRCVYRRGRRLCYWGGR | 64 | >128 | -1.09 | 18 | 1 | 7.91 |
| BHS-35 | RCRCVCGRRKRRCYCGGK | 128 | >128 | -1.33 | 18 | 2 | 7.82 |
| BHS-36 | RGRCVCRRRKCRCYWGRT | 4 | 64 | -1.2 | 18 | 2 | 7.85 |
| BHS-37 | RCKCVYIRRRRRCFCRGR | 32 | >128 | -1.59 | 18 | 2 | 8.85 |
| BHS-38 | RGRCVCGRRRCRCYYRRR | 8 | >128 | -0.96 | 18 | 2 | 8.85 |
| BHS-39 | RGRCVCRRRKCRCFWRGK | 32 | 128 | -1.29 | 18 | 2 | 8.85 |
| BHS-40 | RCKCVCRRRRRRCFCGGK | 128 | >128 | -1.81 | 18 | 2 | 8.82 |
| BHS-41 | RCRCVCRRRRCRCYCRGT | 4 | 64 | -1.35 | 18 | 3 | 7.79 |

MBC: minimum bactercidal information, MH: Meuller-Hinton, HS: human serum, L2FC: $log_2$-fold change in reads, #S-S: number of disulfide bonds found in the majority of molecular population

**Table S4: SAP serum optimization library.**

| Name | Sequence | MH MBC (µg/ml) | HS MBC (µg/ml) | %Hemo |
|---|---|---|---|---|
| SAP | RCRCVCRRRKCRCYCGRT | 4 | 64 | nd |
| SAP-1 | RCRCVCRRRKCLCQCRRT | 4 | 64 | 0.1 ± 0.2 |
| SAP-2 | RCRCVCRRRKCLCQCRRT$^{A}$ | 4 | 32 | 0.2 ± 0.1 |
| SAP-3 | RCRCVCRRRKCLCQCRR- | 16 | 128 | -0.1 ± 0.0 |
| SAP-4 | -CRCVCRRRKCLCQCRRT | 4 | 128 | 6.4 ± 0.5 |
| SAP-5 | -CRCVCRRRKCLCQCRR- | 16 | >128 | 0.1 ± 0.2 |
| SAP-6 | RCRCVCRRRKCLCQCR-- | 4 | 64 | 0.2 ± 0.1 |
| SAP-7 | -CRCVCRRRKCLCQC--- | 4 | 128 | 0.2 ± 0.1 |
| SAP-8 | RCRCVCRRRKCLCQCRR-$^{A}$ | 8 | 32 | 1.7 ± 0.1 |
| SAP-9 | -CRCVCRRRKCLCQCRRT$^{A}$ | 32 | >128 | -0.1 ± 0.2 |
| SAP-10 | -CRCVCRRRKCLCQCRR-$^{A}$ | 4 | 64 | 0 ± 0.1 |
| SAP-11 | RCRCVCRRRKCLCQCR--$^{A}$ | 2 | 16 | 0.5 ± 0.0 |
| SAP-12 | -CRCVCRRRKCLCQC---$^{A}$ | 2 | 64 | -0.1 ± 0.1 |
| SAP-13 | RCRCVCRRRKCLCQCRRT | 8 | 128 | 0.2 ± 0.0 |
| SAP-14 | RCRCVCRRRKCLCQCRRT | 4 | 128 | 29.1 ± 0.5 |
| SAP-15 | RCRCVCRRRKCLCQCRRT | >32 | >128 | -0.1 ± 0.0 |
| SAP-16 | dCRCVCRRRKCLCQCddT | 2 | 64 | 0 ± 0.1 |
| SAP-17 | RCRCVCddddCLCQCRRT | 2 | 32 | -0.1 ± 0.0 |
| SAP-18 | dCdCVCddddCLCQCddT | 4 | 32 | 0 ± 0.0 |
| SAP-19 | oCRCVCRRRKCLCQCooT | 2 | 64 | -0.1 ± 0.1 |
| SAP-20 | RCRCVCooooCLCQCRRT | 2 | 32 | 0 ± 0.1 |
| SAP-21 | oCoCVCooooCLCQCooT | 8 | 128 | 0 ± 0.0 |
| SAP-22 | pCRCVCRRRKCLCQCppT | 4 | 64 | 0 ± 0.1 |
| SAP-23 | RCRCVCppppCLCQCRRT | 8 | >128 | 0 ± 0.0 |
| SAP-24 | pCpCVCppppCLCQCppT | 4 | 128 | -0.2 ± 0.1 |
| SAP-25 | $^{OcA}$RCRCVCRRRKCLCQCR--$^{A}$ | 16 | 64 | 7.2 ± 0.4 |
| SAP-26 | RCRCVCddddCLCQCR--$^{A}$ | 2 | 16 | -0.2 ± 0.0 |
| SAP-27 | RCRCVCRRRKCLCQCR-- | >32 | >128 | nd |

MIC: minimum inhibitory concentration, MBC: minimum bactercidal information, MH: Meuller-Hinton, HS: human serum, %Hemo: percent hemolysis at 128 µg/ml, error is one standard deviation of triplicate samples, $^{A}$: amidation, Underline: D-enantiomer, d: diaminobutyric acid, o: ornithine, p: diamino proprionic acid, $^{OcA}$: octanoic acid

**Table S5. SAP-26 spectrum of antibacterial activity**

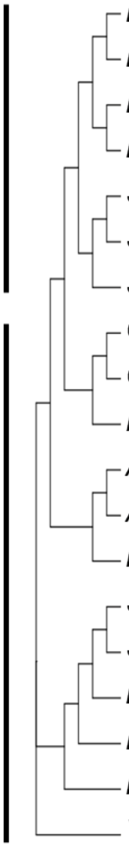

| Group | Bacterial strain | SAP-26 MH MIC (µg/ml) |
|---|---|---|
| Monoderm | *Enterococcus faecium* AR01 | 32 |
| Monoderm | *Listeria monocytogenes* ATCC BAA-679 | 2 |
| Monoderm | *Bacillus cereus* ATCC 14579 | 32 |
| Monoderm | *Bacillus subtilis* PY79 | 1 |
| Monoderm | *Staphylococcus aureus* USA100 | 32 |
| Monoderm | *Staphylococcus aureus* ATCC 43300 | 32 |
| Monoderm | *Staphylococcus epidermidus* ATCC 12228 | 4 |
| Diderm | *Corynebacterium psuedodiphtheriticum* ATCC 10700 | 4 |
| Diderm | *Corynebacterium striatum* ATCC 6940 | 0.5 |
| Diderm | *Mycobacterium smegmatis* ATCC 700084 | 32 |
| Diderm | *Acinetobacter baumannii* AB5075 | 64 |
| Diderm | *Acinetobacter baumannii* ATCC AYE | 16 |
| Diderm | *Pseudomonas aeruginosa* ATCC 27853 | 8 |
| Diderm | *Salmonella typhimurium* LT2 | 4 |
| Diderm | *Shigella flexneria* SA100 | 4 |
| Diderm | ***Escherichia coli* ATCC 25922** | 2 |
| Diderm | *Enterobacter cloacae* ATCC 13047 | >64 |
| Diderm | *Klebsiella pneumoniae* MKP103 | >64 |
| Diderm | *Vibrio cholerae* C6706 | 8 |

MH = Mueller-Hinton; MIC = minimum inhibitory concentration

**Table S6. Plasmids and Oligonucleotides.**

| Plasmids | Source |
|---|---|
| pMMBEH67_lpp_ompA | (16) |
| pDM1_empty | (32) |
| pDM1_mcr-1 | (32) |
| pUltraGFP | (38) |
| **Oligonucleotides** | **Sequence** |
| oJR557 - F BH library | gtattggtaccagtcaagagcctg |
| oJR598 - R BH library | ctg cag gtc gac tta TBT TCV TCY MYA GWA GCA TMG GCR THT TCY CCT TMY AYA AAC GCA TYT GCV CCT ggt tcc tcc gat acc cgc ag |
| 2x(NR)tether gBlock | ATTGCCGATGGTACACGTCAAGTCAAGAGCCTGCAGCGCCCGCCGCAGAGGCGACTCCTGCTGCTGAAGCTCCAGCTAGCGAAGCGCCTGCAGCAGAAGCTGCCCCAGCGGATGCTGCCGAAGCCCCAGCCGCTGGCATCAGTCAGGAACCTGCTGCACCAGCTGCGGAAGCTACACCAGCAGCGGAGGCACCAGCGAGTGAAGCACCGGCTGCGGAAGCCGCTCCTGCAGATGCCGCTGAGGCTCCAGCTGCGGGTATCGGAGGAACCCGCGGTGGGCGTCTTTGTTA |
| F amplicon | aatgATACGGCGACCACCGAGATCTACACTCTTTCCCTACACGACGCTCTTCCGATCTCTCCAGCTGCGGGTATCGGAGGA |
| R index 1 | CAAGCAGAAGACGGCATACGAGATCGTGATGTGACTGGAGTTCAGACGTGTGCTCTTCCGATCTgccaagcttgcatgcctgcaggtcgacTTA |
| R index 2 | CAAGCAGAAGACGGCATACGAGATACATCGGTGACTGGAGTTCAGACGTGTGCTCTTCCGATCTgccaagcttgcatgcctgcaggtcgacTTA |
| R index 3 | CAAGCAGAAGACGGCATACGAGATGCCTAAGTGACTGGAGTTCAGACGTGTGCTCTTCCGATCTgccaagcttgcatgcctgcaggtcgacTTA |
| R index 4 | CAAGCAGAAGACGGCATACGAGATTGGTCAGTGACTGGAGTTCAGACGTGTGCTCTTCCGATCTgccaagcttgcatgcctgcaggtcgacTTA |
| R index 5 | CAAGCAGAAGACGGCATACGAGATCACTGTGTGACTGGAGTTCAGACGTGTGCTCTTCCGATCTgccaagcttgcatgcctgcaggtcgacTTA |
| R index 6 | CAAGCAGAAGACGGCATACGAGATATTGGCGTGACTGGAGTTCAGACGTGTGCTCTTCCGATCTgccaagcttgcatgcctgcaggtcgacTTA |
| R index 7 | CAAGCAGAAGACGGCATACGAGATGATCTGGTGACTGGAGTTCAGACGTGTGCTCTTCCGATCTgccaagcttgcatgcctgcaggtcgacTTA |
| R index 8 | CAAGCAGAAGACGGCATACGAGATTCAAGTGTGACTGGAGTTCAGACGTGTGCTCTTCCGATCTgccaagcttgcatgcctgcaggtcgacTTA |
| R index 9 | CAAGCAGAAGACGGCATACGAGATCTGATCGTGACTGGAGTTCAGACGTGTGCTCTTCCGATCTgccaagcttgcatgcctgcaggtcgacTTA |
| R index 10 | CAAGCAGAAGACGGCATACGAGATAAGCTAGTGACTGGAGTTCAGACGTGTGCTCTTCCGATCTgccaagcttgcatgcctgcaggtcgacTTA |
| R index 11 | CAAGCAGAAGACGGCATACGAGATGTAGCCGTGACTGGAGTTCAGACGTGTGCTCTTCCGATCTgccaagcttgcatgcctgcaggtcgacTTA |
| R index 12 | CAAGCAGAAGACGGCATACGAGATTACAAGGTGACTGGAGTTCAGACGTGTGCTCTTCCGATCTgccaagcttgcatgcctgcaggtcgacTTA |

All oligonucleotides were ordered from Integrated DNA technologies (IDT). IDT single letter nucleotide base notations are used.

**Table S7: Strains used.**

| Strains | Source |
|---|---|
| *E. coli* W3110 | Lab Stock |
| *E. coli* 25922 | Lab Stock |
| *Enterococcus faecium AR01* | Lab Stock |
| *Listeria monocytogenes ATCC BAA-679* | Lab Stock |
| *Bacillus cereus ATCC 14579* | Lab Stock |
| *Bacillus subtilis PY79* | Lab Stock |
| *Staphylococcus aureus USA100* | Lab Stock |
| *Staphylococcus aureus ATCC 43300* | Lab Stock |
| *Staphylococcus epidermidus ATCC 12228* | Lab Stock |
| *Corynebacterium psuedodiphtheriticum ATCC 10700* | Lab Stock |
| *Corynebacterium striatum ATCC 6940* | Lab Stock |
| *Mycobacterium smegmatis ATCC 700084* | Lab Stock |
| *Acinetobacter baumannii AB5075* | Lab Stock |
| *Pseudomonas aeruginosa ATCC 27853* | Lab Stock |
| *Salmonella typhimurium LT2* | Lab Stock |
| *Shigella flexneria SA100* | Lab Stock |
| *Escherichia coli ATCC 25922* | Lab Stock |
| *Enterobacter cloacae ATCC 13047* | Lab Stock |
| *Klebsiella pneumoniae MKP103* | Lab Stock |
| *Vibrio cholerae C6706* | Lab Stock |
